# Sex-specific acute cerebrovascular response to photothrombotic stroke in mice requires rho-kinase

**DOI:** 10.1101/2022.10.07.511301

**Authors:** Joanna Raman-Nair, Gregory Cron, Kathleen McLeod, Baptiste Lacoste

## Abstract

With high energy consumption and low energy storage, the brain is highly reliant on continuous cerebral blood flow (CBF) that delivers substrates to maintain proper function, which is compromised after a stroke. The current study explores the overlapping roles played by two important modulators of cerebrovascular tone, rho-kinase (ROCK) and endogenous sex hormones, in the acute CBF responses to a photothrombotic (PT) model of ischemic stroke in *ROCK2*^*+/-*^ mice and wild-type (WT) littermates. To remove endogenous hormones, male mice were gonadectomized and female mice were ovariectomized, whereas control (“intact”) animals received a sham surgery prior to stroke induction. Intact WT males showed a delayed drop in CBF compared to intact WT females, where maximal CBF drop was observed 48 hours following stroke. Gonadectomy in males did not alter this response, however ovariectomy in females produced a “male-like” response. Intact *ROCK2*^*+/-*^ males also showed such phenotypic response, which was not altered by gonadectomy. Alternatively, intact *ROCK2*^*+/-*^ females showed a striking difference in CBF values compared to intact WT females, where they displayed higher CBF values immediately post-stroke and also showed a maximal CBF drop 48 hours post-stroke, which was not altered by ovariectomy. Overall, there is a marked sex difference in acute CBF responses to PT stroke, which appears to be mediated by endogenous female sex hormones and ROCK2. This study reveals important sex-differences and the involvement of ROCK2 in acute CBF responses to PT stroke in mice.

**Significance Statement:** There are very few mechanistic investigations on disparities between sexes in post-stroke CBF outcome. Rho-kinase, an important regulator of vascular tone, has only been explored in males in terms of its modulation of CBF following stroke. Both rho-kinase and endogenous female sex hormones have a converging role on the regulation of endothelial nitric oxide synthase (eNOS), an important modulator of vascular tone. Rho-kinase is thought to elicit its neuroprotective effects against ischemic stroke through eNOS, however this has never been investigated in both sexes. Elucidating the cellular and molecular bases of sex differences in cerebrovascular pathophysiology is vital for understanding the origins of stroke outcomes, and for designing novel therapeutic strategies to promote functional recovery in both women and men.

## Introduction

By limiting tissue perfusion, stroke affects brain homeostasis, vascular function, and compromises neuronal health (Iadecola & Anrather, 2011; Tymianski, 2011; Moskowitz, et al., 2010). The two major classifications of stroke are ischemic and hemorrhagic, of which ischemic stroke accounts for approximately 85% of all events (Heart & Stroke Foundation of Canada, 2019). Ischemic stroke results from the narrowing or occlusion of a cerebral blood vessel, resulting in immediate local blood flow restriction to the brain. This is extremely detrimental for brain functionality because, unlike other organs, the brain has a limited capacity to store energy and relies on a continuous supply of substrates from blood flow (Willie, et al., 2014). If blood flow is not restored, substantial cell death occurs in the core of the injury which can not be salvaged, resulting in debilitating functional consequences to the patient (Heiss, 2012; Hossmann, 2012).

Biological sex markedly influences cerebral blood flow (CBF) as well as the prevalence and progression of cardiovascular diseases, including stroke (Cowan, et al., 2017; Cosgrove, et al., 2007; Krause, et al., 2006; Orshal & Khalil, 2004). Epidemiological studies show that stroke incidence dramatically increases in women following menopause (Bonkhoff, et al., 2021; Wang, et al., 2019; Barker-Collo, et al., 2015; Appelros, et al., 2009). Because women have longer average life expectancies, post-menopausal women have higher rates of stroke compared to age-matched men, resulting in increased mortality, worse psychological outcomes, and higher rates of disability (Heart & Stroke Foundation of Canada, 2019; Madsen, et al., 2019; Ahnstedt, et al., 2016; Persky, et al., 2010; Turtzo & McCullough, 2010). Following menopause, estrogen production by the ovaries decreases drastically which has led to the presumption that estrogen is protective against cardiovascular disease. This is thought to be mediated by the upregulation of endothelial nitric oxide synthase (eNOS) via estrogen signaling (Boese, et al., 2017; Novella, et al., 2012; Miyazaki-Akita, et al., 2007; Simoncini, et al., 2000). Increased activation and transcription of eNOS by estrogens ultimately results in increased bioavailability of nitric oxide (NO), which is a potent vasodilator of vascular smooth muscle cells (VSMCs) and thereby modulates vascular tone (Chen, et al., 2008). For this reason, estrogen is said to reduce systemic blood pressure, protect against vascular disease, and increase CBF to the brain (Turtzo & McCullough, 2008).

Removal of endogenous female sex hormones through ovariectomy (Ovx) has been shown to decrease CBF following experimental ischemic stroke in rodents compared to females with their ovaries left intact, however supplementation with estrogen to restore blood flow has shown conflicting results (McCullough, et al., 2001; Yang, et al., 2000). Only one study has shown that CBF values of female rats are higher following experimental stroke compared to males and Ovx females (Alkayed, et al., 1998), and the mechanism behind this difference has yet to be investigated. Female rats have also shown to have quicker vascular remodeling of occluded and peripheral vessels compared to males following ischemic stroke (Yang, et al., 2019). Overall, specific knowledge on sex differences in cerebrovascular disease is limited, and the regulation of endothelial function by sex hormones in disease states is poorly understood.

Another important regulator of endothelial function and vascular tone is rho-associated coiled-coil containing protein kinase (ROCK), which is activated by the upstream effector RhoA. RhoA/ROCK signaling serves many roles, including regulation of cell contractility through inhibitory action on myosin light chain phosphatase (MLCP) by ROCK2, the isoform expressed primarily in the brain and vasculature (Çiçek & Ayaz, 2015; Hartmann, et al., 2015; Wang, et al., 2009). Upregulation of ROCK leads to increased vascular contraction and calcium sensitization of VSMCs (Nunes & Webb, 2021). ROCK2 has therefore been implicated in the pathogenesis of various vascular diseases, including hypertension (Hartmann, et al., 2015; Abd-Elrahman, et al., 2015; Rankinen, et al., 2008), age-related vascular dysfunction (Nunes & Webb, 2021), vascular dysfunction associated with diabetes (Çiçek & Ayaz, 2015; Soliman, et al., 2012), and stroke (Hiroi, et al., 2018). Moreover, in both peripheral (Ahnstedt, et al., 2013; Nuno, et al., 2009; Nuno, et al., 2007; Lamping & Faraci, 2001) and cerebral vessels (Faraci, et al., 2006; Chrissobolis, et al., 2004), ROCK has been shown to be implicated in sex differences in cerebrovascular reactivity.

ROCK also influences vascular tone by directly inhibiting both the activation (Sugimoto, et al., 2007; Ming, et al., 2002) and expression of eNOS (Laufs & Liao, 1998). Furthermore, RhoA/ROCK signaling has been shown to be upregulated in human endothelial cells during hypoxia, mediating downregulation of eNOS expression and activation (Jin, et al., 2006; Takemoto, et al., 2002). ROCK activity has been shown to be upregulated following ischemic stroke, contributing to increased vascular permeability and enhancing oxidative stress (Cui, et al., 2013; Allen, et al., 2010; Satoh, et al., 2010; Shin, et al., 2007). Specifically pertaining to the ROCK2 isoform, selective ROCK2 inhibition after experimental ischemic stroke dose-dependently reduced infarct volume and limited perfusion loss in male mice (Lee, et al., 2014). Male *ROCK2*^*+/-*^ mice, who exhibit constitutively enhanced eNOS expression in brain endothelial cells, also have reduced infarct volumes following experimental ischemic stroke (Hiroi, et al., 2018). These neuroprotective effects correlated with higher levels of NO, and neuroprotective effects were abolished in eNOS deficient (*eNOS*^*-/-*^) mice (Hiroi, et al., 2018). Thus, ROCK signaling during ischemia may contribute to vasoconstriction and reduced blood flow in the hypoxic region by increasing VSMC contractility, which in turn may be mediated by the eNOS/NO pathway.

The mechanisms underlying cerebrovascular stroke outcomes are poorly understood, and the effects of sex hormones on cerebrovascular regulation in the ischemic brain have yet to be fully comprehended. Considering the overlapping roles of hormonal and rho-kinase regulation of vascular function, this study aims to address the overarching hypothesis that ROCK2 is involved in CBF outcomes following a focal ischemic stroke in a sex-specific manner.

## Methods

### Subjects

Male and female *ROCK2*^*+/-*^ mice and wild-type (WT) littermates were bred in house and housed a maximum of five per cage with free access to food and water. Animals were maintained on Teklad Global 18% Protein Rodent Diet (Harlan Laboratories, Teklad Diets, Madison WI) composed of 18.6% protein, 6.2% fat, 3.5% fiber and 44.2% carbohydrates. Mice were aged 8-10 weeks when experimental procedures were initiated. All methods and procedures were approved by the University of Ottawa’s Animal Care Committee and are in accordance with the Canadian Council on Animal Care guidelines.

### Female Gonadectomy

All female mice received either a sham surgery (referred to herein as “intact females”) or a bilateral ovariectomy (Ovx). Mice were initially anesthetized with 4% isoflurane and then maintained at 2.5% isoflurane for the duration of the surgery. The back of the mouse was shaved and skin was disinfected with 70% isopropyl alcohol and 4% chlorhexidine. Slow-release buprenorphine (1.2mg/kg, Chiron Compounding Pharmacy Inc., Guelph, ON, Canada) and 1mL of saline (0.9%) were both administered subcutaneously (SC) prior to performing the surgery. The mouse was then placed on a pad heated to 37°C and positioned in sternal recumbency. Using a sterile scalpel, a 1-cm incision was made down the midline of the lower back. The skin was then gently separated using a blunt probe and the ovary was visualized near the flanks, where it is attached to adipose tissue. A small incision was made in the abdominal wall, and the adipose tissue was gently pulled out of the intraperitoneal cavity along with the ovary attached. A clamp was placed just below the ovary at the uterine horn and held in place for 5-10 seconds, followed by excision of the ovary. The clamp remained in place for an additional 5-10 seconds after excision to reduce bleeding. Once the clamp was released, the adipose tissue was returned to the peritoneal cavity and the abdominal incision was closed with 6-0 prolene sutures. This exact procedure was performed again on the opposite side to remove both ovaries. Finally, the midline incision in the back was closed with autoclips and topical bupivacaine (bupivacaine hydrochloride as monohydrate, 2%, Chiron Compounding Pharmacy Inc., Guelph, ON, Canada) was applied to the incision site for analgesia. Mice were placed in a 37°C chamber until they woke up and were then returned to a clean home cage. Mice were monitored 4 hours following the procedure and for the following 3 days in the morning to ensure proper recovery. Sham Ovx surgery was performed exactly as above, without clamping or excision of the ovaries. Mice were allowed to recover for 10-14 days before any further experimental procedures were performed.

### Male Gonadectomy

All male mice received either a sham surgery (referred to herein as “intact males”) or gonadectomy (Gdx) to remove both testicles. Mice were initially anesthetized with 4% isoflurane and then maintained at 2.5% isoflurane for the duration of the surgery. The scrotum of the mouse was shaved and skin was disinfected with 70% isopropyl alcohol and 4% chlorhexidine. Carprofen (20mg/kg, Rimadyl®, Zoetis Canada Inc., Kirkland, QC, Canada) and 1mL of saline (0.9%) were both administered SC prior to performing the surgery. The mouse was then placed on a pad heated to 37°C and positioned in dorsal recumbency. A 1-cm incision was made down the midline of the scrotum and the tunica was gently separated from the skin using a blunt probe. Gentle pressure was applied to the lower abdomen of the mouse to push out both testes. A clamp was then placed on the adipose tissue visible just above the testes to restrict blood flow. After approximately 10 seconds, a size 4-0 suture was tied loosely around the tissue behind the clamp. The clamp was then released, the knot was checked to ensure no skin was caught, and then the knot was tied tightly around the tissue. The testes were clamped one more time in front of the knotted suture and then both testicles were excised. The clamp remained in place for another 5-10 seconds. After removing the clamp, the excision site was monitored for any bleeding and then the incision site was closed with 6-0 prolene sutures. Topical bupivacaine (2%) was applied to the incision site for analgesia and mice were placed in a 37°C chamber until they woke up and were then returned to the home cage. Mice were monitored for 4 hours following the procedure and for the following 2 days in the morning to ensure proper recovery. Mice also received a second SC dose of carprofen (20mg/kg) the morning following the procedure for analgesia. Sham Gdx surgery was performed exactly as above, without clamping, tying off, or excision of the testes. Mice were allowed to recover for 10-14 days before any further experimental procedures were performed.

### Laser Doppler Flowmetry and Photothrombotic Stroke

Cerebral blood flow was measured using laser doppler flowmetry (LDF, transonic® tissue perfusion monitor, Ithaca, NY, USA) in the somatosensory cortex under ketamine (100mg/kg, Vétoquinol N.-A. Inc., Lavaltrie, QC, Canada) and xylazine (10mg/kg, Nerfasin 20™, Dechra Regulatory B. V., Bladel, The Netherlands) anesthesia (K/X) administered SC as a bolus dose. A top up dose of ketamine (25mg/kg) and xylazine (2.5mg/kg) was administered SC as a maintenance dose approximately 30 minutes following the initial dose. Animal temperature was maintained with a feedback-controlled rectal thermometer and heating pad system (Harvard Apparatus Homeothermic Monitoring System, Holliston, MA, USA) at 37°C ± 1°C. After a sufficient anesthetic plane was reached, the mouse was placed in a stereotaxic frame and an incision was made down the midline of the skull. A high-speed microdrill was then used to thin the skull to translucency at the following coordinates relative to bregma: -2.7 anterior-posterior, +4 medial-lateral. At these coordinates, an LDF probe was placed just above the skull at a 25° angle to measure blood flow. The LDF technique uses a probe comprised of a laser of a specific monochromatic wavelength and a detector. The laser is reflected off red blood cells and the backscattered light creates an interference pattern on the detector surface, which measures the doppler shift in frequency the backscattered light as red blood cells pass, providing a relative measure of perfusion (Fredriksson, et al., 2007). While LDF does not give an absolute measurement of perfusion, it has high temporal resolution to measure rapid relative changes in perfusion, which can be measured reliably over time (Tajima, et al., 2014). To induce photothrombotic (PT) stroke, mice received an intraperitoneal injection of the photosensitive dye rose bengal (RB, 100mg/kg, MilliporeSigma: Cat. No. R3877) dissolved in phosphate buffered saline (PBS). RB was allowed to circulate for 5 minutes, during which baseline LDF measurements were taken. At the end of the 5-minute period, a 532nm laser was turned on for 10 minutes at a distance of 3-cm from the skull at the same coordinates used for LDF measurements. Photoactivation of the light-sensitive RB results in production of singlet oxygen leading to endothelium damage and platelet aggregation forming a thrombus (Lee et al., 2007). Post-stroke LDF measurements were then taken at the same coordinates for 30 minutes. Post-stroke blood flow measurements were normalized to baseline for each individual mouse for statistical analysis. At the end of the procedure, mice received a dose of buprenorphine (0.1mg/kg, Vetergesic® buprenorphine hydrochloride, Ceva Animal Health Inc., Cambridge, ON, Canada) administered SC for analgesia and 1.5mL of SC saline (0.9%) to rehydrate the animal. Finally, topical bupivacaine (2%) was applied to the incision site for analgesia and mice were left in a 30°C incubator for approximately 4 hours until the anesthesia had worn off and normal activity resumed.

### Follow up Laser Doppler Flowmetry Measurements

Additional 30-minute LDF measurements were taken at 48-hours and 1-week post-stroke. Mice were anesthetized with ketamine (100mg/kg) and xylazine (10mg/kg) anesthesia, with no maintenance dose being required. Mice were again placed in a stereotaxic frame and an LDF probe was placed at a 25° angle at the same coordinates used for all previous LDF measurements and PT stroke induction: -2.7 anterior-posterior, +4 medial-lateral. No additional thinning of the skull was performed. Animal temperature was maintained with a rectal thermometer and heating pad at 37°C ± 1°C. At the end of the procedure, mice received 1.5mL of SC saline (0.9%), topical bupivacaine (2%) was applied to the incision site, and mice were left in a 30°C incubator to recover for 4 hours.

### Infarct Volumes

For quantification of infarct volumes, in-vivo magnetic resonance imaging (MRI) was performed 48 hours after PT stroke induction using a 7 Tesla GE/Agilent MR (University of Ottawa pre-clinical imaging core facility). Mice were anesthetized for the MRI procedure using 2% isoflurane. A 2D fast spin echo sequence (FSE) pulse sequence was used for the imaging, with the following parameters: slice thickness=0.5 mm, spacing=0 mm, field of view=2.5 cm, matrix=256×256, echo time=41 ms, repetition time=7000 ms, echo train length=8, bandwidth=16 kHz, fat saturation. Stroke lesions demonstrated hyperintensity. MRI images were loaded in the Fiji software (https://imagej.net/software/fiji/) and infarct volumes were quantified using a custom script for outlining the infarct perimeter. All infarct volumes were quantified twice to obtain an average of the two measures for each animal. Subsequent LDF measurements for this timepoint were taken at least 1 hour following isoflurane exposure.

### Nitric Oxide Staining

PT stroke was performed as outlined above. 24 hours following stroke induction, mice were sacrificed using cervical dislocation followed by decapitation and brains were extracted and placed in ice-cold 1xPBS. A brain matrix placed on ice was used to cut 2-mm sections of tissue that encompassed the infarcted tissue. Brain slices were incubated in the dark with 5μmol of DAF-FM diacetate stain (Invitrogen by ThermoFisher, cat# D23844) diluted in 1xPBS for 20 minutes followed by a 1xPBS wash with DAPI [1:20 000] for 5 minutes. A final 1xPBS wash was performed for at least 10 minutes prior to being imaged. Just before imaging, the slice was removed from solution and placed on a microscope slide for imaging using a Zeiss Axio Imager M2 microscope equipped with a digital camera (Axiocam 506 mono) at 5X objective. DAPI was excited with 461nm and DAF-FM diacetate was excited using 594nm wavelength. Using Fiji software (https://imagej.net/software/fiji/), infarcted tissue was outlined with the polygon tool on the DAPI channel only. This outline was then saved as a region of interest (ROI) and loaded directly onto the same image with only the DAF-FM stain and the mean grey value for the entire ROI was calculated by the software. The ROI was then positioned on the same brain slice on the hemisphere contralateral to the lesion and in a corresponding area of the cortex and the mean grey value for the DAF-FM stain was also taken. Mean grey value from the ROI on the contralateral side was subtracted from the mean grey value of the infarcted tissue and this is the value that was used for analysis.

### Statistical Analysis

LDF measurements provided continuous CBF measurements over the selected region. To analyze post-stroke CBF values, measurements were normalized to pre-stroke baseline values after RB was injected, but before laser illumination. To do this, the percent change in CBF was measured by taking each post-stroke data point and dividing it by the mean of the entire 4-minute baseline recording, and multiplying that value by 100%. Each baseline data point was also divided by the mean of the entire 4-minute baseline recording and multiplied by 100%. Therefore, normalized pre-stroke baseline values are always 100%, and post-stroke values measure the change in CBF from baseline values. Statistical analyses were performed using GraphPad Prism software (version 9.2.0). Hyperacute timepoints were analyzed using a repeated measures two-way ANOVA. Due to animal loss, comparisons of 48-hour and 1 week data points were analyzed using a two-way ANOVA.

## Results

### Sex Differences in CBF Following a PT Stroke in the Somatosensory Cortex

Following PT stroke in the somatosensory cortex, CBF averaged over the 30-minute period dropped to 85.13% ± 2.48 (mean ± SD) of baseline values in Intact WT males, while Intact WT females dropped to 62.78% ± 2.75. There was no significant difference between these groups in overall average CBF values in the first 30 minutes post stroke (**Fig. 1C**). Although there was no significant difference in CBF values between overall CBF values post-stroke, the individual curves for each group (**Fig. 1A**) show a phenotypic difference between groups. Intact WT females show an immediate reduction in CBF and stay relatively stable throughout the 30-minute recording, whereas Intact WT males show a noticeably sloped delay in CBF drop. Because of this apparent separation between groups at the beginning of the CBF values post-stroke, we broke up the 30-minute time period into smaller sections, herein referred to as “hyperacute timepoints”. The most noticeable difference between groups was during the first 0-5 minutes following stroke, where Intact WT males had mean normalized CBF values of 88.60% ± 27.73 and Intact WT females averaged 59.34% ± 19.47, however there was no statistical significance between groups (*p*=0.0635). There were no differences between groups at any other hyperacute timepoint (**Fig. 1B**). Interestingly, when measured at 48 hours following stroke, Intact WT males showed a significant decrease in CBF values compared to the values taken immediately post-stroke (**Fig. 1B**). A further drop in CBF values was not observed in Intact WT females, and there were no differences within or between groups at the 1-week timepoint (**Fig. 1B**).

**Figure 1.**
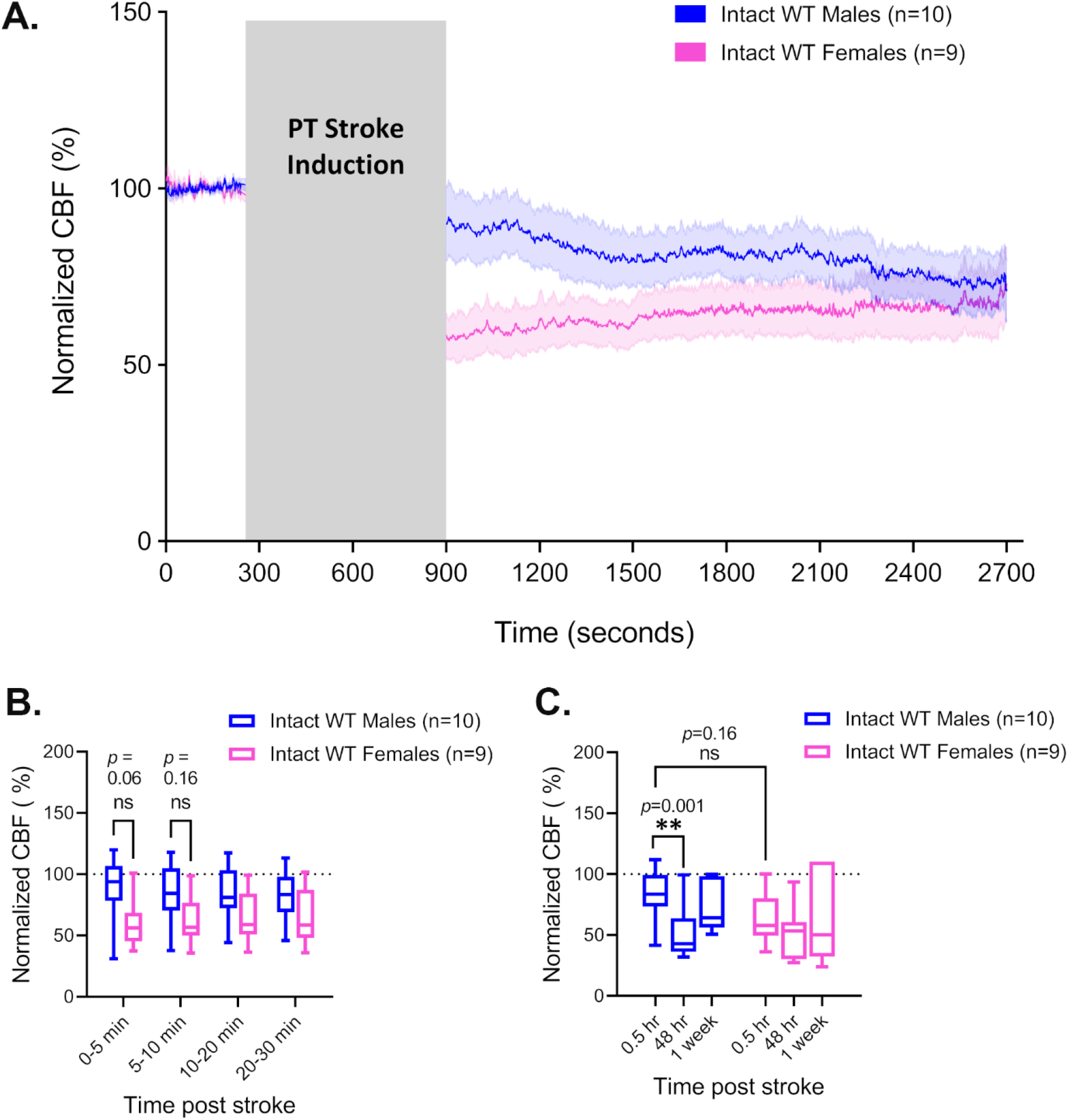
CBF in intact WT males vs. females post PT stroke in the somatosensory cortex. (**A**) CBF measured by LDF under K/X anesthesia in intact WT male and intact WT female mice before and after a PT stroke. Grey box indicates time passed during laser irradiation of the PT stroke induction. Post-PT values are normalized to pre-PT baseline values. Curved lines represent average normalized CBF of all animals in the respective group. Shaded area above and below curves represent SEM. (**B**) Averaged CBF measurements of hyperacute timepoints during the 30 minutes immediately following PT stroke induction shown in A. Bars represent min to max values with a line at the mean ± SD. (**C**) Averaged 30-minute recordings of normalized CBF measured immediately post stroke (0.5hr), 48 hours post PT, and 1 week post PT. Bars represent min to max values with a line at the mean ± SD. \*\**p*<0.01 (2-way ANOVA and Sidak post-hoc tests)

### The Contribution of Endogenous Sex Hormones to Observed Sex Differences in CBF Following PT Stroke

To determine the role of endogenous sex hormones on CBF outcomes following stroke, mice were gonadectomized prior to receiving a PT stroke, and CBF values were compared to intact mice that received a control sham surgery. Gdx WT male mice did not show any differences in CBF values compared to Intact WT males at any hyperacute timepoint following stroke (**Fig. 2A,B**). Furthermore, as observed in Intact WT males, Gdx WT males also showed a significant decrease in CBF values at 48 hours post-stroke when compared to immediate post-stroke values (**Fig. 2C**). There were no differences in CBF values within or between groups at the 1-week timepoint (**Fig. 2C**).

**Figure 2.**
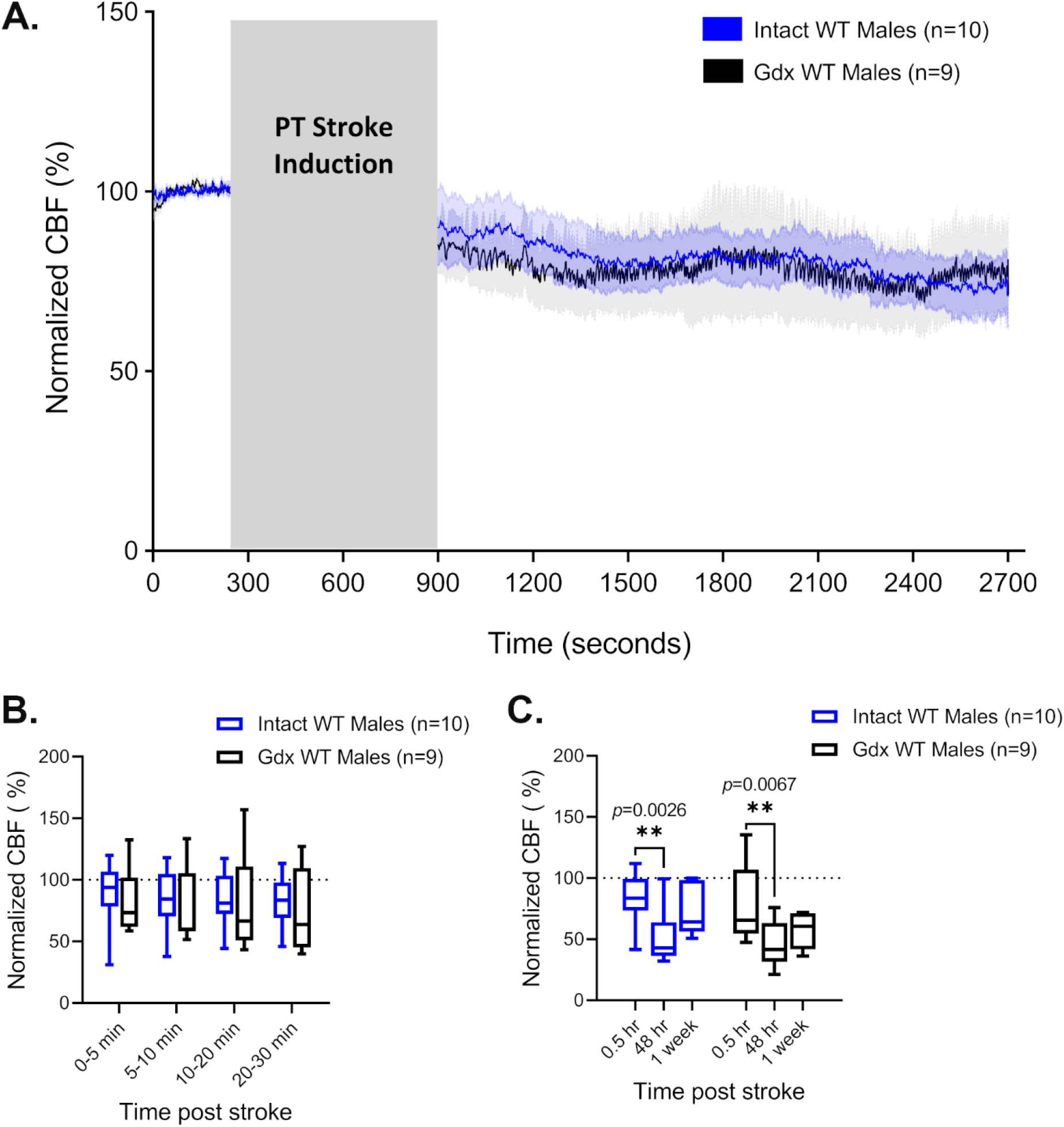
Removing endogenous male sex hormones does not change CBF outcomes post PT stroke. (**A**) CBF measured by LDF under K/X anesthesia in Intact WT male and Gdx WT male mice before and after a PT stroke. Grey box indicates time passed during laser irradiation of the PT stroke induction. Post-PT values are normalized to pre-PT baseline values. Curved lines represent average normalized CBF of all animals in the respective group. Shaded area above and below curves represent SEM. (**B**) Averaged CBF measurements of hyperacute timepoints during the 30 minutes immediately following PT stroke induction shown in A. Bars represent min to max values with a line at the mean ± SD. (**C**) Averaged 30-minute recordings of normalized CBF measured immediately post stroke (0.5hr), 48 hours post PT, and 1 week post PT. Bars represent min to max values with a line at the mean ± SD. \*\**p*<0.01 (2-way ANOVA and Sidak post-hoc tests)

CBF values in Ovx WT females dropped to an average of 80.22% ± 20.64 compared to baseline values during the full 30-minute post-stroke period. This is higher than was observed in Intact WT females, although the difference was not statistically significant (**Fig. 3C**). Although there is separation between the CBF curves immediately post-stroke (**Fig. 3A**), there was no significant difference in CBF values between groups at any of the hyperacute timepoints (**Fig. 3B**). Interestingly, similar to what was observed in Intact and Gdx WT males, Ovx WT females also showed a significant decrease in CBF values measured at 48 hours post-stroke compared to those measured immediately post-stroke (**Fig. 3C**). There were no differences in CBF values within or between groups at the 1-week timepoint (**Fig. 3C**).

**Figure 3.**
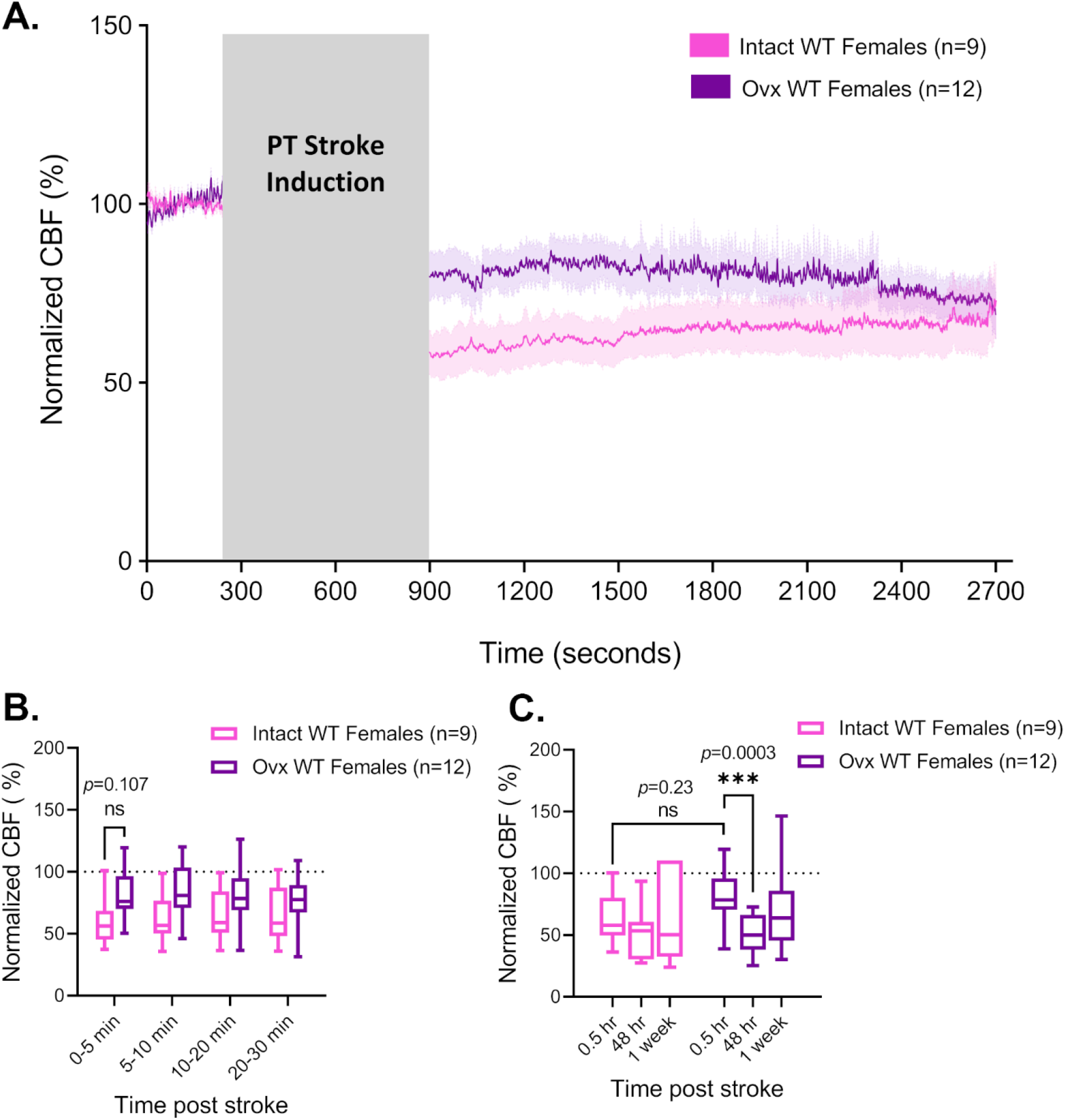
The effects of removing endogenous female sex hormones on CBF outcomes following stroke. (**A**) CBF measured by LDF under K/X anesthesia in Intact WT female and Ovx WT female mice before and after a PT stroke. Grey box indicates time passed during laser irradiation of the PT stroke induction. Post-PT values are normalized to pre-PT baseline values. Curved lines represent average normalized CBF of all animals in the respective group. Shaded area above and below curves represent SEM. (**B**) Averaged CBF measurements of hyperacute timepoints during the 30 minutes immediately following PT stroke induction shown in A. Bars represent min to max values with a line at the mean ± SD. (**C**) Averaged 30-minute recordings of normalized CBF measured immediately post stroke (0.5hr), 48 hours post PT, and 1 week post PT. Bars represent min to max values with a line at the mean ± SD. \*\*\**p*<0.001 (2-way ANOVA and Sidak post-hoc tests)

### Sex-specific Outcomes of ROCK2 Haploinsufficiency on CBF Values Following PT stroke

Intact *ROCK2*^*+/-*^ male mice did not show any differences in CBF values compared to Intact WT males at any hyperacute timepoint following stroke (**Fig. 4A,B**). Furthermore, as observed in Intact WT males, Intact *ROCK2*^*+/-*^ males also showed a significant decrease in CBF values at 48 hours post-stroke when compared to immediate post-stroke values (**Fig. 4C**). There were no differences in CBF values within or between groups at the 1-week timepoint (**Fig. 4C**). This suggests that ROCK2 haploinsufficiency in males does not alter CBF outcomes following a PT stroke in the somatosensory cortex.

**Figure 4.**
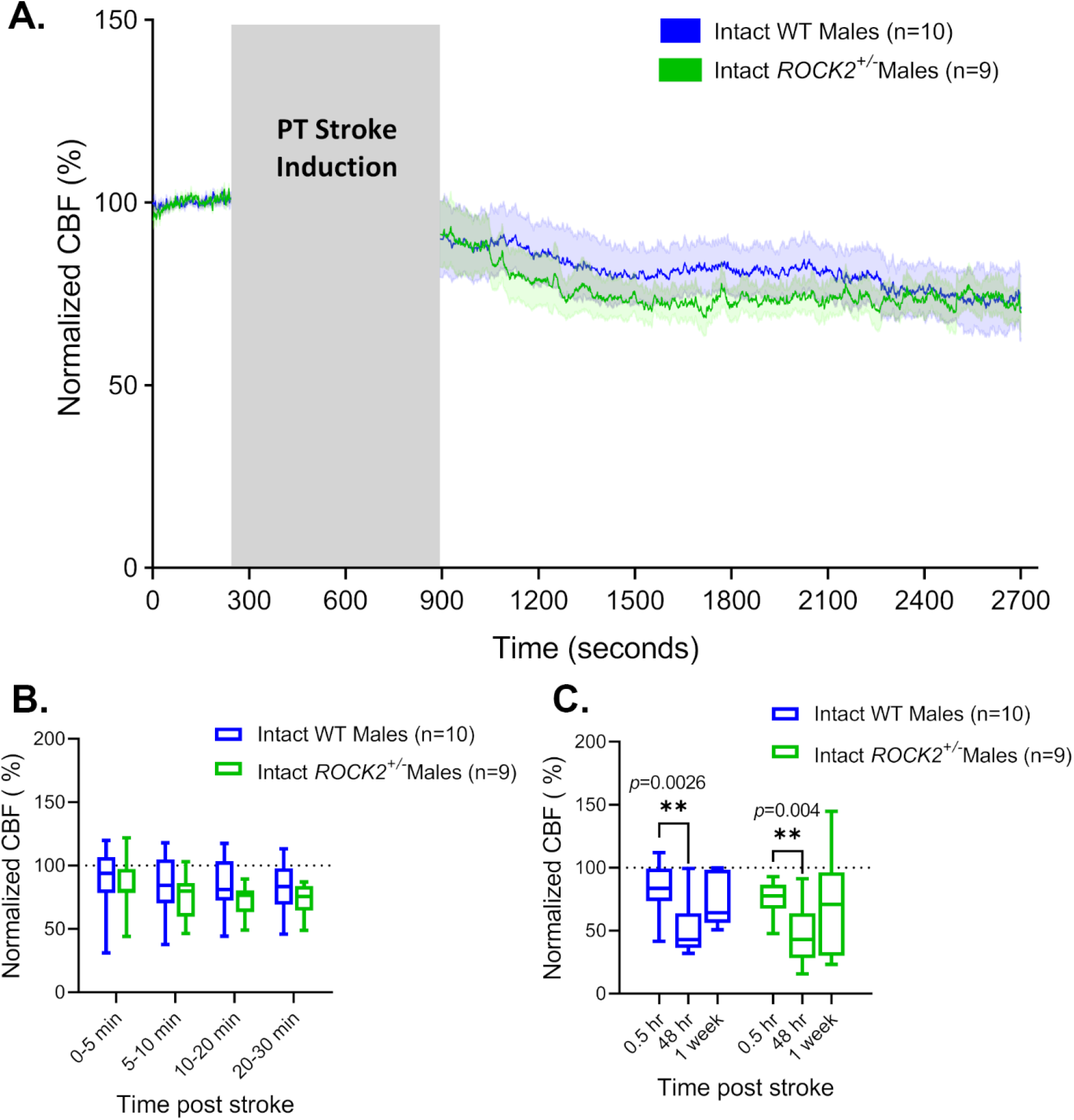
ROCK2 haploinsufficiency does not alter CBF outcomes in males following PT stroke in the somatosensory cortex. (**A**) CBF measured by LDF under K/X anesthesia in Intact WT male and Intact *ROCK2*^*+/-*^ male mice before and after a PT stroke. Grey box indicates time passed during laser irradiation of the PT stroke induction. Post-PT values are normalized to pre-PT baseline values. Curved lines represent average normalized CBF of all animals in the respective group. Shaded area above and below curves represent SEM. (**B**) Averaged CBF measurements of hyperacute timepoints during the 30 minutes immediately following PT stroke induction shown in A. Bars represent min to max values with a line at the mean ± SD. (**C**) Averaged 30-minute recordings of normalized CBF measured immediately post stroke (0.5hr), 48 hours post PT, and 1 week post PT. Bars represent min to max values with a line at the mean ± SD. \*\**p*<0.01 (2-way ANOVA and Sidak post-hoc tests)

Intact *ROCK2*^*+/-*^ females had significantly higher CBF values immediately following stroke compared to Intact WT females when averages for the entire 30-minute timepoint were compared (**Fig. 5A,C**). Surprisingly, when this timepoint was broken up into smaller hyperacute timepoints, there was no difference between groups at any of the hyperacute timepoints (**Fig. 5B**). Intact *ROCK2*^*+/-*^ females also showed a significant decrease in CBF values at 48 hours post-stroke when compared to immediate post-stroke values (**Fig. 5C**). There were no differences in CBF values within or between groups at the 1-week timepoint (**Fig. 5C**).

**Figure 5.**
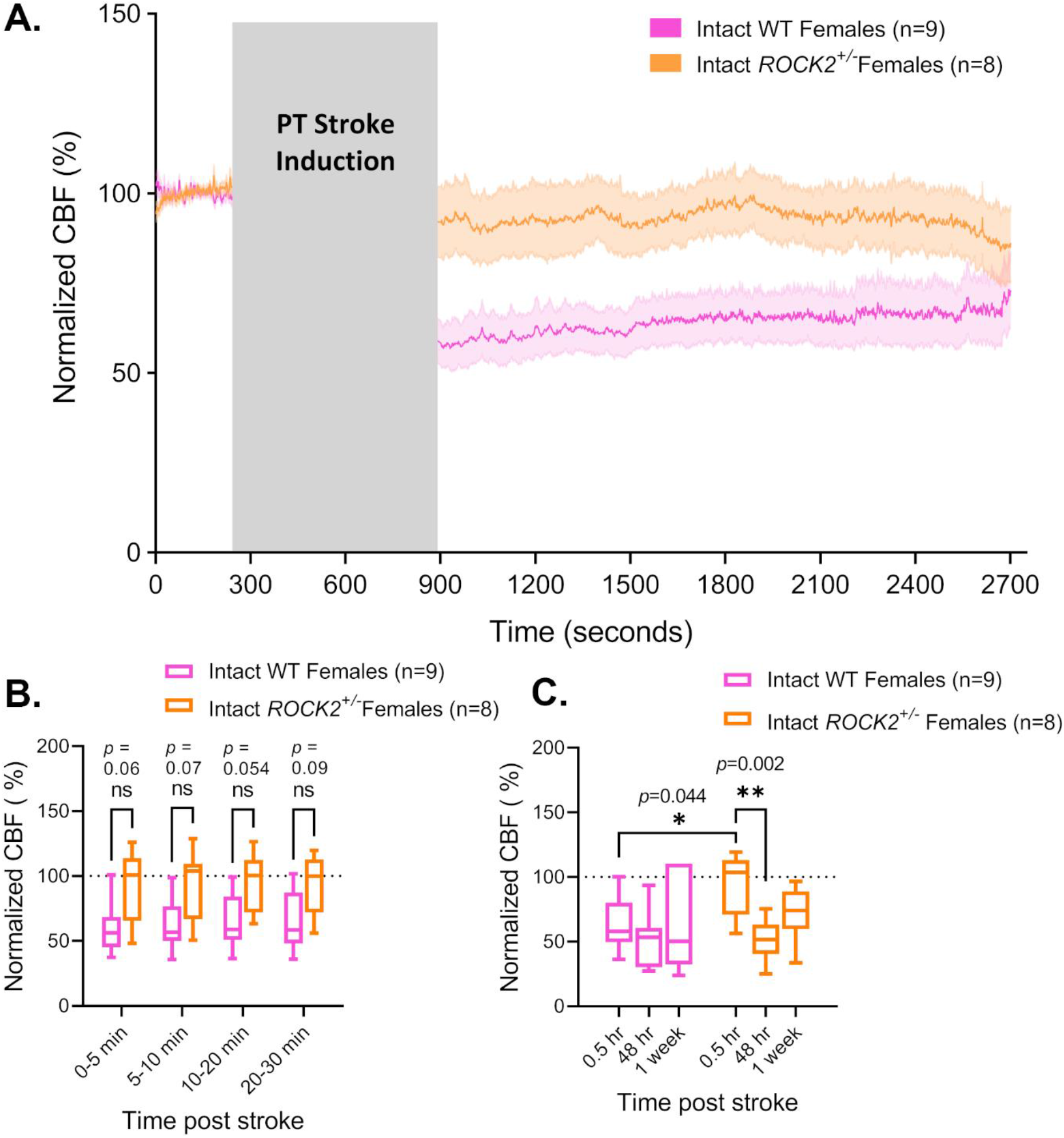
The effects of ROCK2 haploinsufficiency on CBF outcomes in females following PT stroke in the somatosensory cortex. (**A**) CBF measured by LDF under K/X anesthesia in Intact WT female and Intact ROCK2^+/-^ female mice before and after a PT stroke. Grey box indicates time passed during laser irradiation of the PT stroke induction. Post-PT values are normalized to pre-PT baseline values. Curved lines represent average normalized CBF of all animals in the respective group. Shaded area above and below curves represent SEM. (**B**) Averaged CBF measurements of hyperacute timepoints during the 30 minutes immediately following PT stroke induction shown in A. Bars represent min to max values with a line at the mean ± SD. (**C**) Averaged 30-minute recordings of normalized CBF measured immediately post stroke (0.5hr), 48 hours post PT, and 1 week post PT. Bars represent min to max values with a line at the mean ± SD. \*\**p*<0.01, \**p*<0.05 (2-way ANOVA and Sidak post-hoc tests)

### Interactions Between Sex Hormones and ROCK2 in CBF Outcomes Following PT Stroke

There were no differences in CBF values in the total averages or in separated hyperacute timepoints between male and female Intact ROCK2^+/-^ mice following PT stroke (**Fig. 6A,B**). Both Intact *ROCK2*^*+/-*^ males and females showed a significant decrease in CBF values at 48-hours post stroke compared to immediate post-stroke values, with no other differences at any other timepoint (**Fig. 6C**). Removal of male endogenous sex hormones did not change the CBF response in the hyperacute phase following stroke in Gdx *ROCK2*^*+/-*^ males compared to Intact *ROCK2*^*+/-*^males (**Fig. 7A,B**). As observed in Intact *ROCK2*^*+/-*^ males, Gdx *ROCK2*^*+/-*^ males also showed a decrease in CBF values measured 48 hours post-stroke compared to those measured immediately post-stroke, although this was not statistically significant (*p*=0.139, **Fig. 7C**). There were no other differences in CBF values between or within groups at the 1-week timepoint (**Fig. 7C**). Removal of endogenous female sex hormones also did not change the CBF response following PT stroke in the hyperacute phase in Ovx *ROCK2*^*+/-*^ females compared to Intact *ROCK2*^*+/-*^ females (**Fig. 8A,B**). Both Intact and Ovx *ROCK2*^*+/-*^ females showed significantly reduced CBF values at 48 hours following PT stroke compared to values measured immediately following PT stroke (**Fig. 8C**). There were no other differences in CBF values between or within groups at the 1-week timepoint (**Fig. 8C**).

**Figure 6.**
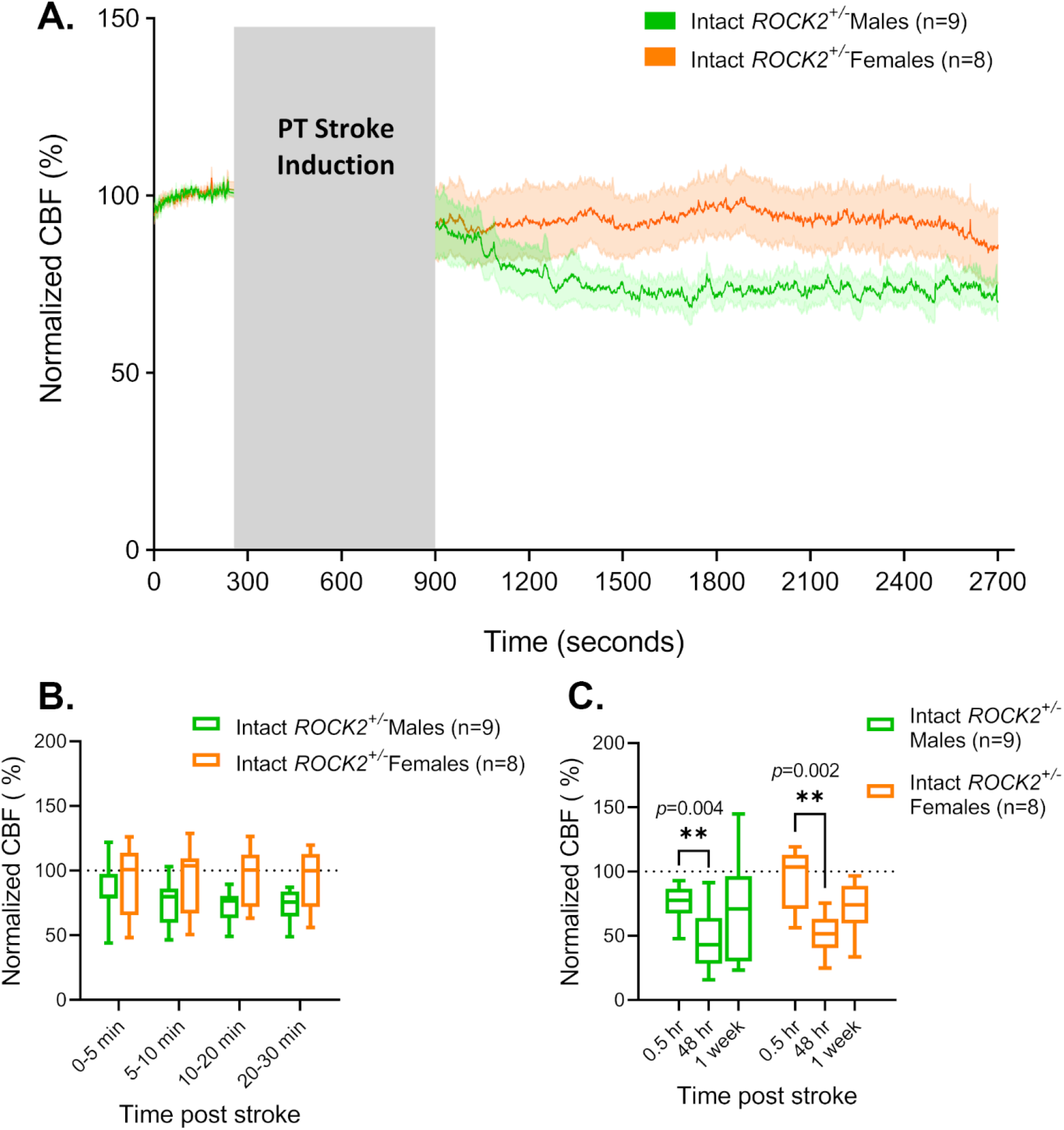
CBF in Intact *ROCK2*^*+/-*^ male vs. female mice post PT stroke in the somatosensory cortex. (A) CBF measured by LDF under K/X anesthesia in Intact *ROCK2*^*+/-*^ male and Intact *ROCK2*^*+/-*^ female mice before and after a PT stroke. Grey box indicates time passed during laser irradiation of the PT stroke induction. Post-PT values are normalized to pre-PT baseline values. Curved lines represent average normalized CBF of all animals in the respective group. Shaded area above and below curves represent SEM. (B) Averaged CBF measurements of hyperacute timepoints during the 30 minutes immediately following PT stroke induction shown in A. Bars represent min to max values with a line at the mean ± SD. (**C**) Averaged 30-minute recordings of normalized CBF measured immediately post stroke (0.5hr), 48 hours post PT, and 1 week post PT. Bars represent min to max values with a line at the mean ± SD. \*\**p*<0.01 (2-way ANOVA and Sidak post-hoc tests)

**Figure 7.**
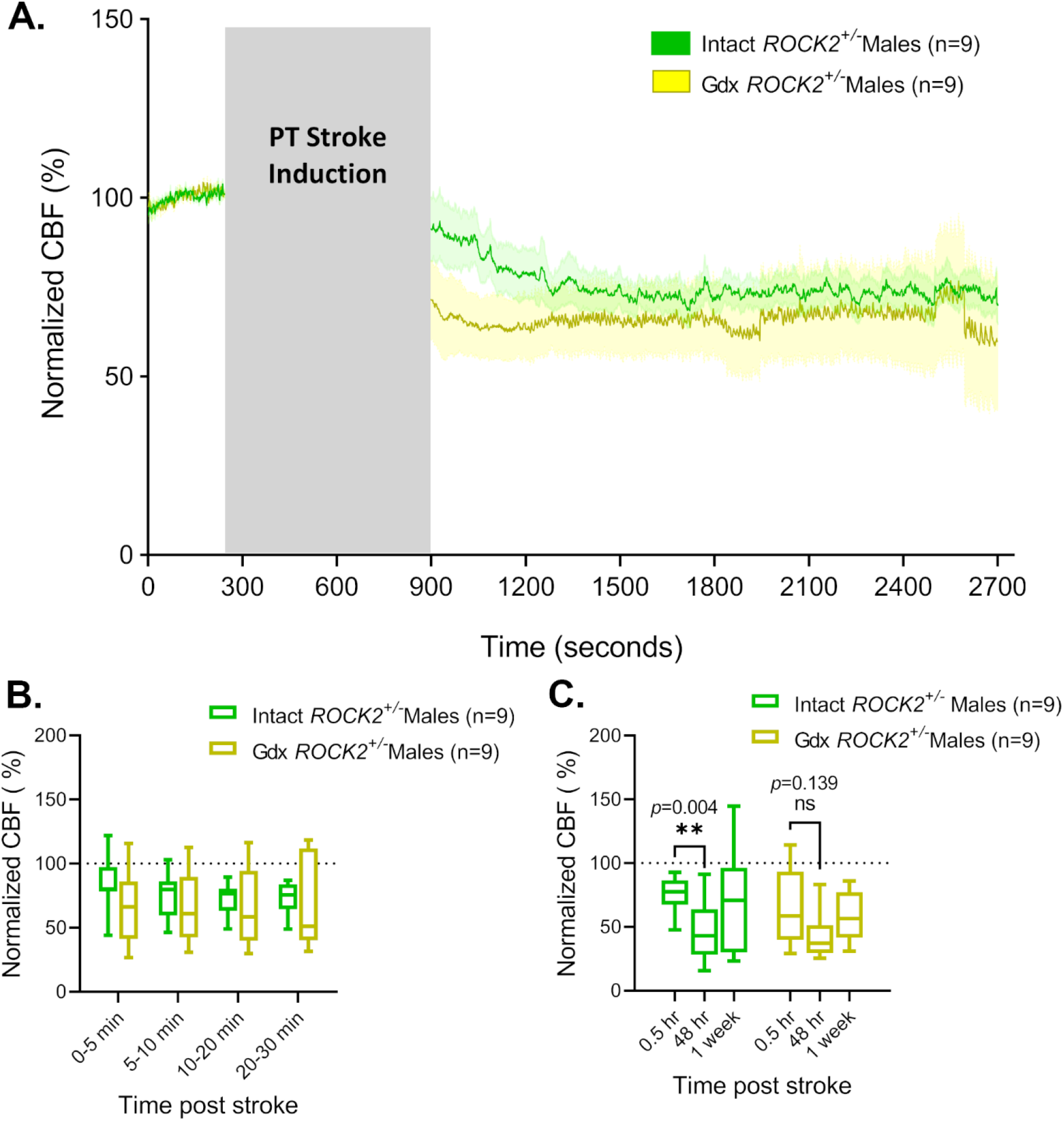
Gonadectomy does not alter the CBF response in *ROCK2*^*+/-*^ males following PT stroke in the somatosensory cortex. (A) CBF measured by LDF under K/X anesthesia in Intact *ROCK2*^*+/-*^ male and Gdx *ROCK2*^*+/-*^ male mice before and after a PT stroke. Grey box indicates time passed during laser irradiation of the PT stroke induction. Post-PT values are normalized to pre-PT baseline values. Curved lines represent average normalized CBF of all animals in the respective group. Shaded area above and below curves represent SEM. (B) Averaged CBF measurements of hyperacute timepoints during the 30 minutes immediately following PT stroke induction shown in A. Bars represent min to max values with a line at the mean ± SD. (**C**) Averaged 30-minute recordings of normalized CBF measured immediately post stroke (0.5hr), 48 hours post PT, and 1 week post PT. Bars represent min to max values with a line at the mean ± SD. \*\**p*<0.01 (2-way ANOVA and Sidak post-hoc tests)

**Figure 8.**
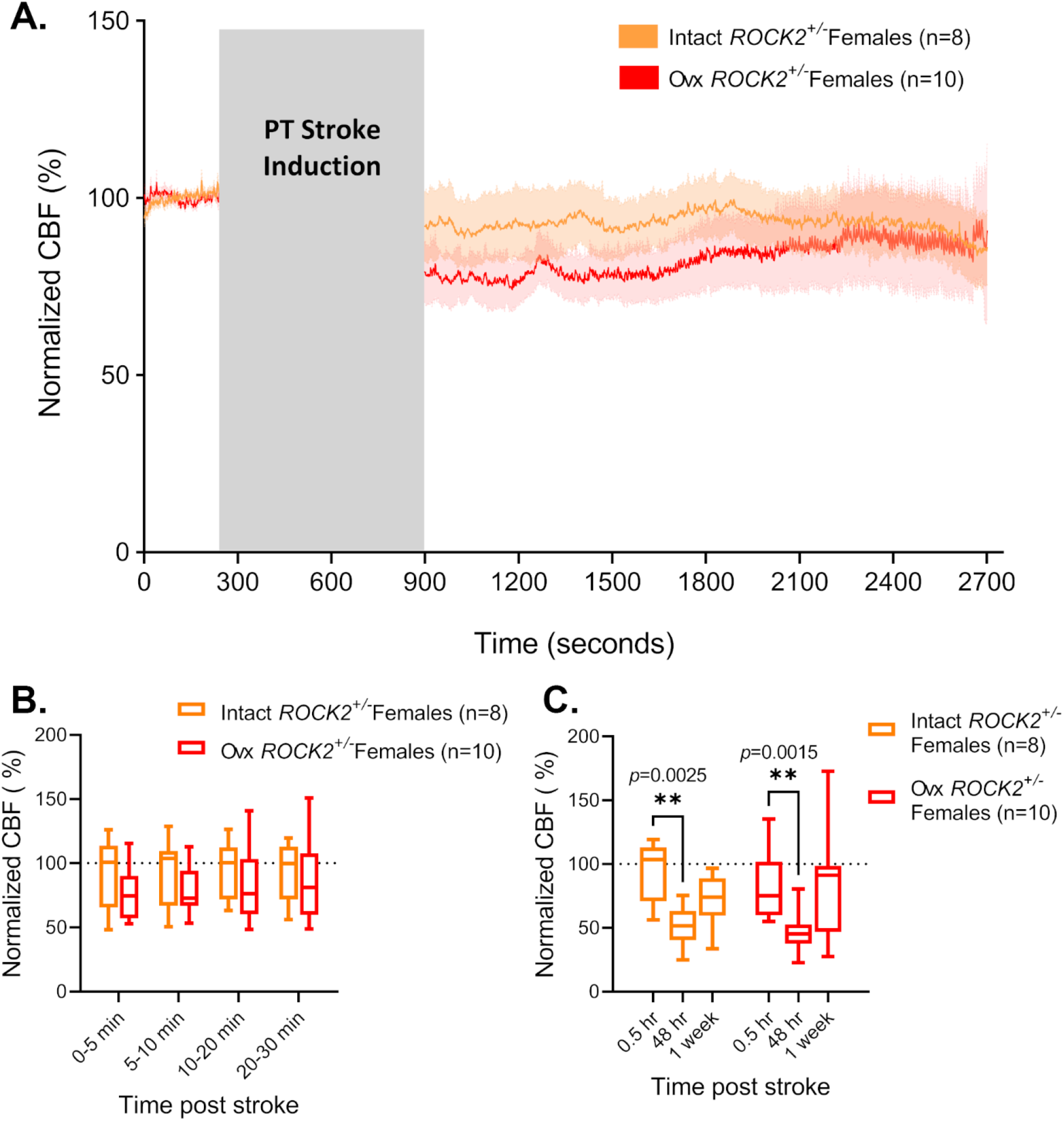
Ovariectomy does not alter the CBF response in ROCK2^+/-^ females following PT stroke in the somatosensory cortex. (A) CBF measured by LDF under K/X anesthesia in Intact *ROCK2*^*+/-*^ female and Ovx *ROCK2*^*+/-*^ female mice before and after a PT stroke. Grey box indicates time passed during laser irradiation of the PT stroke induction. Post-PT values are normalized to pre-PT baseline values. Curved lines represent average normalized CBF of all animals in the respective group. Shaded area above and below curves represent SEM. (B) Averaged CBF measurements of hyperacute timepoints during the 30 minutes immediately following PT stroke induction shown in A. Bars represent min to max values with a line at the mean ± SD. (**C**) Averaged 30-minute recordings of normalized CBF measured immediately post stroke (0.5hr), 48 hours post PT, and 1 week post PT. Bars represent min to max values with a line at the mean ± SD. \*\**p*<0.01 (2-way ANOVA and Sidak post-hoc tests)

Contrary to what was observed in WT mice, removal of endogenous hormones prior to stroke induction in *ROCK2*^*+/-*^ mice did not alter CBF outcomes in either males (**Fig. 9**) or females (**Fig. 10**). Gdx *ROCK2*^*+/-*^ males displayed the same gradual decline in the first 0-5 minutes post stroke as Intact *ROCK2*^*+/-*^ males (**Fig. 9A**), and there were no differences between groups at any hyperacute timepoint (**Fig. 9B**). Gdx *ROCK2*^*+/-*^ males also displayed the same delayed drop in CBF values, reaching maximum CBF deficit when measured at 48 hours following stroke induction, although not statistically significant when compared to immediate post-stroke values (**Fig. 9C**). When compared to Intact *ROCK2*^*+/-*^ females, there were no differences in CBF values in the hyperacute phase in Ovx *ROCK2*^*+/-*^ females (**Fig. 10A,B**). Both groups also showed a maximum drop in CBF at 48 hours post-PT which was significantly different from immediate post-PT values (**Fig. 10C**). There were no other differences in CBF values between or within groups at the 1-week timepoint (**Fig. 10C**).

**Figure 9.**
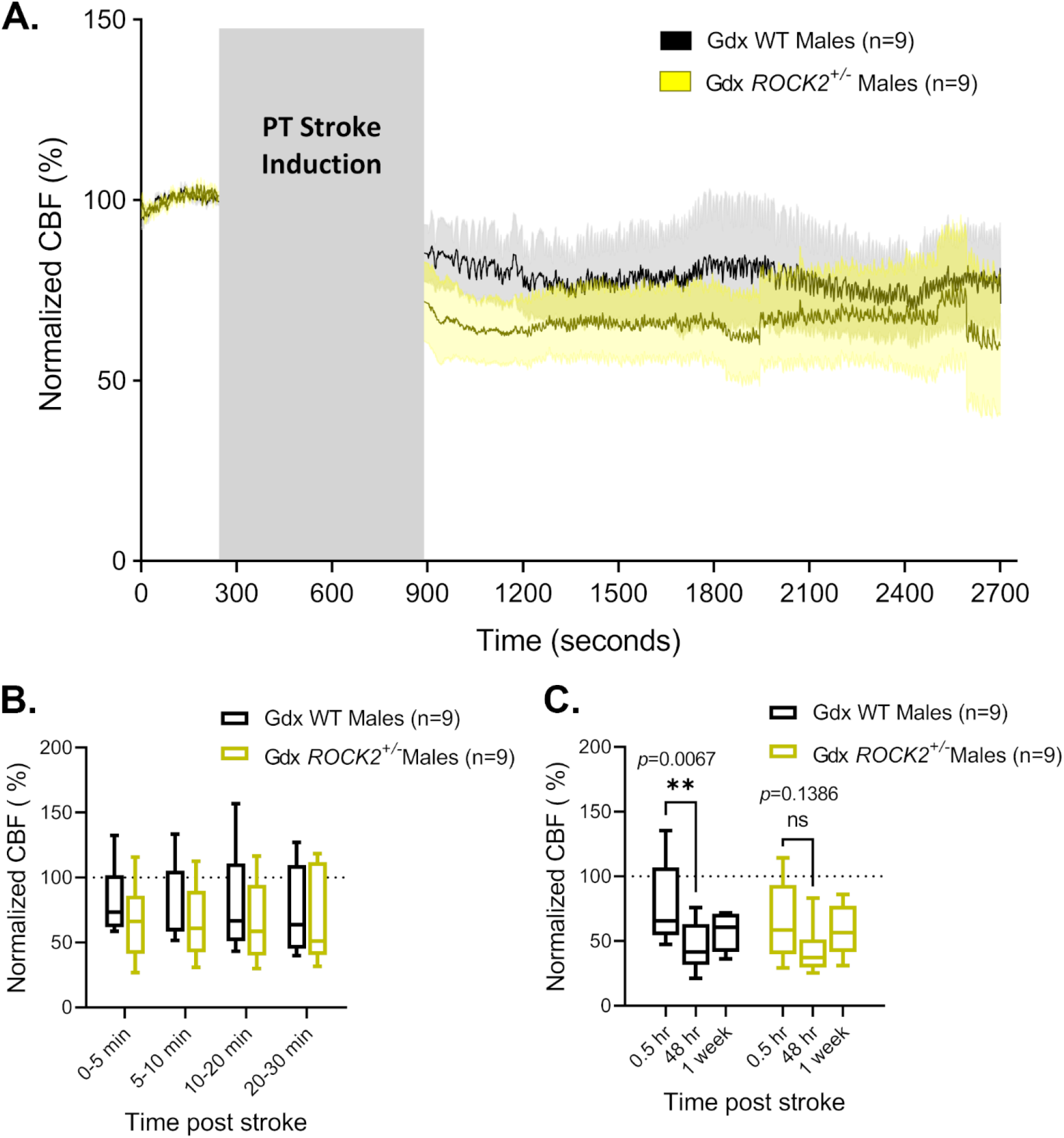
CBF in Gdx WT vs. *ROCK2*^*+/-*^ males post PT stroke in the somatosensory cortex. (**A**) CBF measured by LDF under K/X anesthesia in Gdx WT male and Gdx *ROCK2*^*+/-*^ male mice before and after a PT stroke. Grey box indicates time passed during laser irradiation of the PT stroke induction. Post-PT values are normalized to pre-PT baseline values. Curved lines represent average normalized CBF of all animals in the respective group. Shaded area above and below curves represent SEM. (**B**) Averaged CBF measurements of hyperacute timepoints during the 30 minutes immediately following PT stroke induction shown in A. Bars represent min to max values with a line at the mean ± SD. (**C**) Averaged 30-minute recordings of normalized CBF measured immediately post stroke (0.5hr), 48 hours post PT, and 1 week post PT. Bars represent min to max values with a line at the mean ± SD. \*\**p*<0.01 (2-way ANOVA and Sidak post-hoc tests)

**Figure 10.**
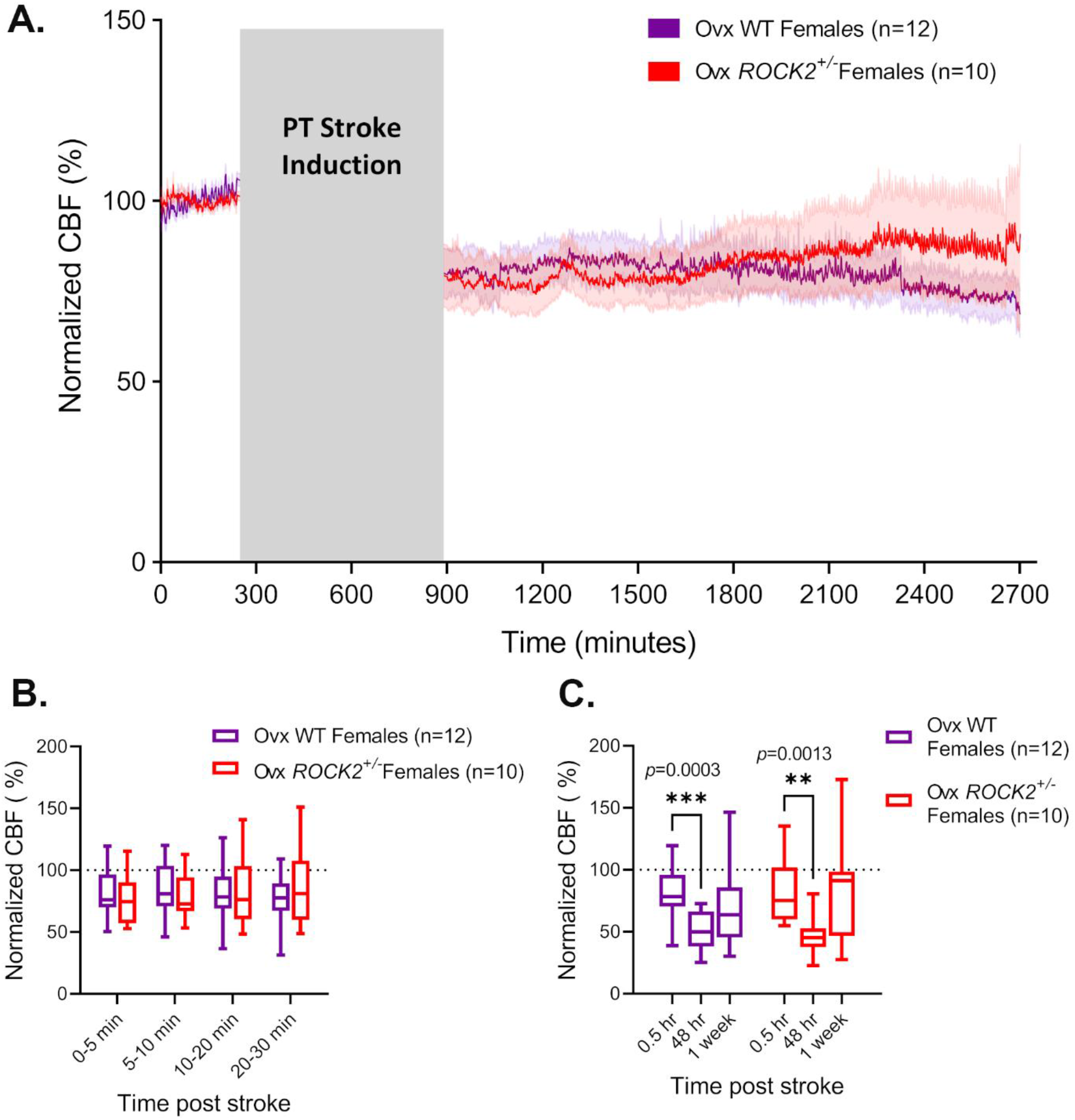
CBF in Ovx WT vs. *ROCK2*^*+/-*^ females post PT stroke in the somatosensory cortex. (**A**) CBF measured by LDF under K/X anesthesia in Ovx WT female and Ovx *ROCK2*^*+/-*^ female mice before and after a PT stroke. Grey box indicates time passed during laser irradiation of the PT stroke induction. Post-PT values are normalized to pre-PT baseline values. Curved lines represent average normalized CBF of all animals in the respective group. Shaded area above and below curves represent SEM. (**B**) Averaged CBF measurements of hyperacute timepoints during the 30 minutes immediately following PT stroke induction shown in A. Bars represent min to max values with a line at the mean ± SD. (**C**) Averaged 30-minute recordings of normalized CBF measured immediately post stroke (0.5hr), 48 hours post PT, and 1 week post PT. Bars represent min to max values with a line at the mean ± SD. \*\*\**p*<0.001 \*\**p*<0.01 (2-way ANOVA and Sidak post-hoc tests)

### Infarct Volume and Nitric Oxide Staining Quantifications

Infarct volumes were measured 48 hours following PT stroke induction from MRI images. A 2-way ANOVA revealed no significant differences in infarct volumes between any of the groups (**Fig. 11A,B**).

**Figure 11.**
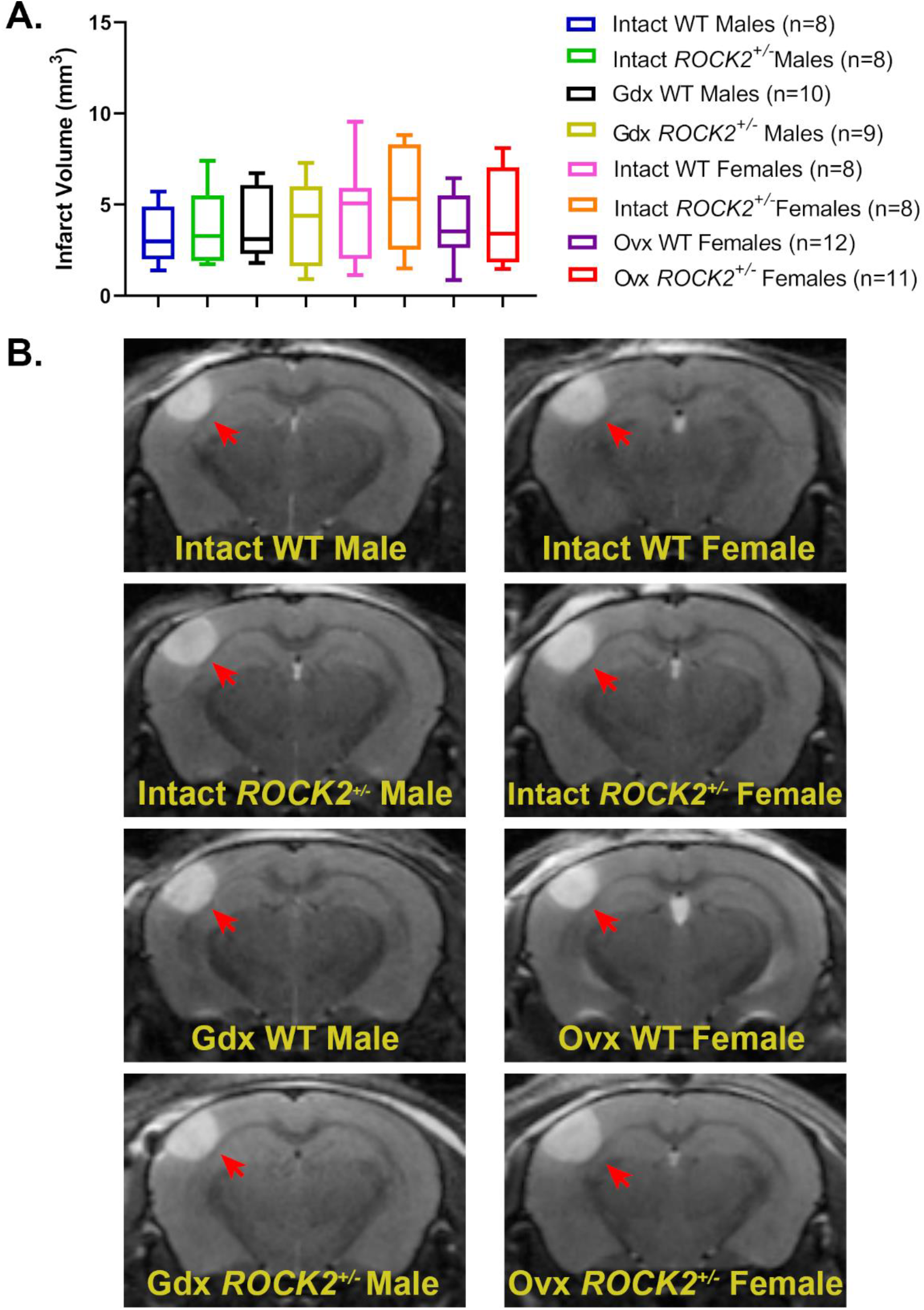
Infarct volume quantification 48 hours following PT stroke. (**A**) Quantification of total infarct volumes measured from magnetic resonance imaging (MRI) images taken 48 hours following PT stroke induction in the somatosensory cortex. Bars represent min to max points with a line at the mean ± SD. (**B**) Representative MRI images of infarcts at 48 hours post PT stroke. Red arrows point to infarct lesion.

Staining for nitric oxide was performed using DAF-FM diacetate and DAPI in 2-mm brain sections 24 hours following PT stroke induction. Average intensity of the DAF-FM stain was quantified in the entire infarcted region, determined by the absence of DAPI staining. A 2-way ANOVA revealed no significant differences in average DAF-FM staining intensity between any of the groups (**Fig. 12A,B**).

**Figure 12.**
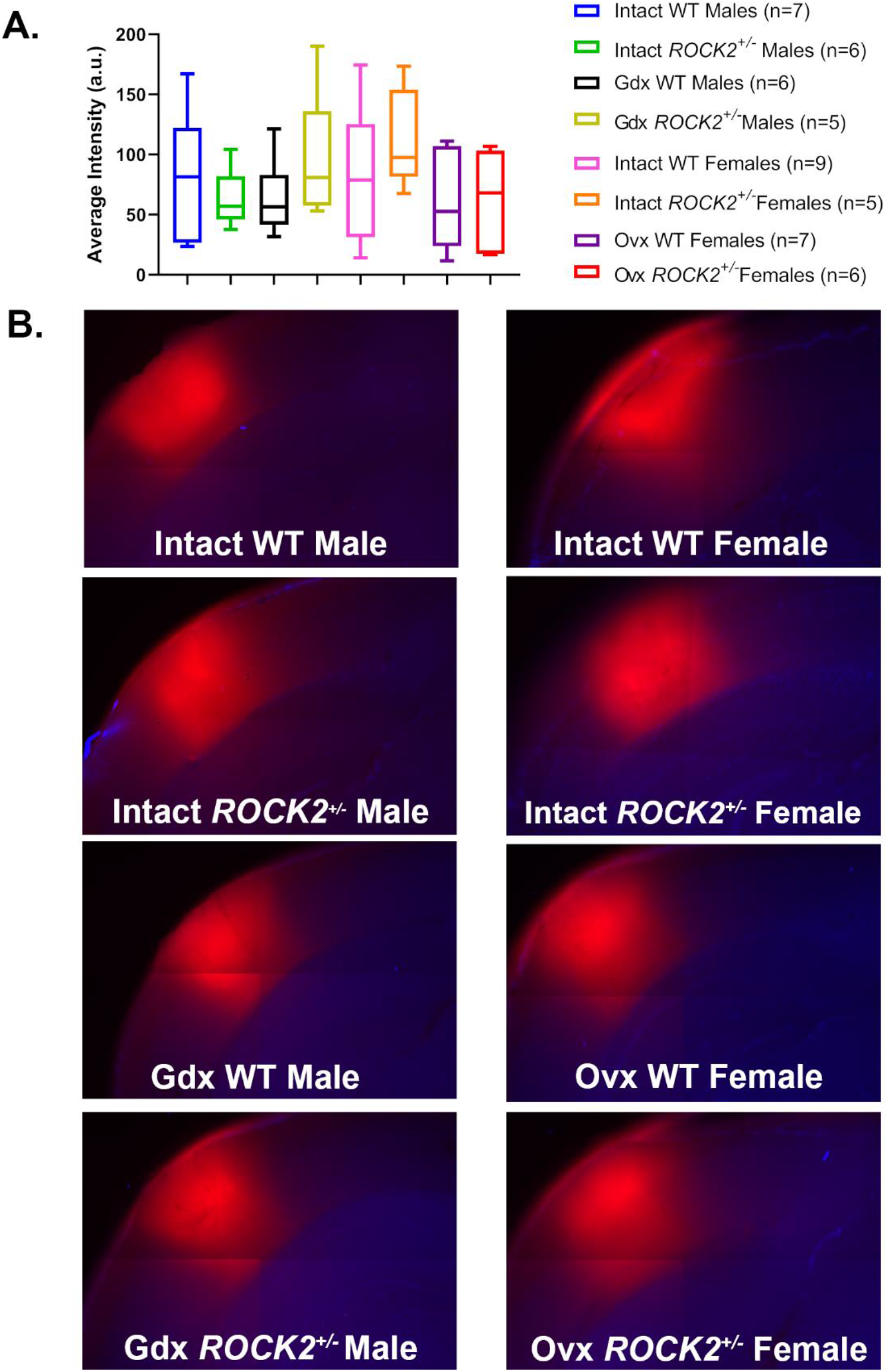
Nitric oxide staining in the infarct 24 hours following PT stroke. (**A**) Quantification of average intensity encompassing entire infarct after staining with DAF-FM diacetate staining performed 24 hours following PT stroke induction in the somatosensory cortex. Bars represent min to max points with a line at the mean ± SD. (**B**) Representative images of brain slices stained with DAF-FM diacetate (red) and DAPI (blue).

## Discussion

The current study investigated CBF outcomes following PT stroke using LDF to provide high temporal resolution of the CBF response. CBF was characterized immediately following stroke and up to one week following stroke in intact and gonadectomized male and female *ROCK2*^*+/-*^ mice and their WT littermates. This study provides novel insight into the mechanisms involved in sex differences in CBF values following ischemia in a preclinical stroke mode. Overall, there is a marked difference between males and females in the acute CBF responses to PT stroke, which appears to be mediated by endogenous female sex hormones and rho-kinase.

### CBF Outcomes in Intact Wild-type Males and Females Following PT Stroke

The CBF response in Intact WT males displayed a phenotypic delayed drop in CBF in the immediate 30-minute timepoint post-stroke compared to Intact WT females. Intact WT males also show a further reduction in CBF at the 48-hour timepoint post stroke, whereas Intact WT females show no change at this timepoint. This apparent sex difference may be due to several factors, including differences in the way each sex responds to stroke induction for this particular stroke model. PT stroke induction involves initiation of the coagulation cascade, resulting in thrombus formation. In both animal models and human studies, females have higher levels of platelets in circulation, and furthermore, platelets from females display higher reactivity and are more prone to aggregation and thrombus formation (Friede, et al., 2020; Miller, et al., 2014; Otahbachi, et al., 2010; Becker, et al., 2006; Leng, et al., 2004; Green, et al., 1992; Zwierzina, et al., 1987). Because Intact WT males appear to take longer to reach maximal CBF deficit – exhibiting peak CBF drop when measured 48 hours following stroke induction – it is possible that PT stroke induction may require more time for platelet aggregation to occur, resulting in the observed delay in CBF drop. Indeed, evidence that female platelets are more reactive and prone to thrombosis supports the idea that thrombus formation resulting from PT stroke induction may have a quicker onset in the female sex.

The only study to directly compare CBF between males and females following ischemic stroke showed that after 2 hours of transient intraluminal middle cerebral artery occlusion (MCAo), intact female rats had higher CBF values measured by LDF under halothane anesthesia compared to both intact males as well as Ovx females (Alkayed, et al., 1998). This improvement in CBF deficit also corresponded to decreased infarct volumes in intact females. There were no improvements at any timepoint post stroke in CBF values in females in the current study, which may be due to mechanistic properties of the stroke model used. Transient MCAo model of ischemic stroke is mechanistically different than PT stroke, involving ligation of the MCA for a pre-determined amount of time followed by rapid reperfusion, and can involve a secondary injury called reperfusion injury. The PT stroke model does not involve any reperfusion, only that which occurs by endogenous fibrinolysis of the thrombotic clot, which results in a much slower and incomplete reperfusion of the affected tissue. Therefore, measuring CBF following MCAo is also assessing the response to reperfusion injury and is a measure of neuroprotection, not just occlusion of the vessel. While females appear to be protected in stroke models involving rapid reperfusion, this may not be the case for the PT stroke model.

### Contribution of Endogenous Sex Hormones to CBF Outcomes Following PT Stroke

Endogenous male sex hormones were removed by gonadectomy of WT males a minimum of 10 days prior to inducing PT stroke and measuring CBF values. There were no differences in the hyperacute phase post stroke of CBF values in Gdx WT males when compared to Intact WT males. Similarly, Gdx WT males showed the same delay in reaching maximal CBF drop post PT as Intact WT males. This suggests that removing endogenous male sex hormones does not alter CBF outcomes following a PT stroke in the somatosensory cortex.

Endogenous female sex hormones were removed by bilateral ovariectomy of WT females a minimum of 10 days prior to PT stroke induction. Interestingly, Ovx of WT females produced a similar phenotype as was observed in both Intact and Gdx WT males. Although not statistically significant, there is a clear separation between the curves of Ovx WT females and Intact WT females in the 0-5 minute hyperacute phase following stroke, where Ovx WT females have slightly higher CBF values immediately following stroke, showing the same phenotype as seen in both aforementioned male groups. Moreover, Ovx WT females also show a delayed drop in CBF, reaching a maximal drop in CBF when measured 48 hours following stroke. These results suggest that endogenous female sex hormones may be involved in the thrombotic response of PT stroke induction. Indeed, chronic E2 treatment exacerbated platelet aggregation in response to endothelial injury of cerebral microvessels in both male and female mice (Rosenblum, et al., 1985). Conversely, chronic testosterone or DHT treatment of male mice increased platelet aggregation following endothelial injury to mesenteric arteries, but not cerebral pial vessels, and furthermore, had no effect on aggregation in female mice (Rosenblum, et al., 1987). Therefore, it is possible that removing endogenous estrogens from female mice through Ovx results in less platelet aggregation and thrombus formation during PT stroke induction, producing this delayed drop in CBF and the same phenotypic response that was observed in males.

### Rho-kinase modulates the CBF response to PT stroke in a sex-specific manner

CBF outcomes were not different at any timepoint in intact males between WT and *ROCK2*^*+/-*^ genotypes, suggesting that ROCK2 haploinsufficiency does not alter the CBF response to stroke in males. While ROCK deletion and inhibition has been shown to be neuroprotective against tissue damage and behavioural outcomes of ischemic stroke, there are somewhat conflicting results in terms of the role of ROCK in CBF outcomes following stroke in preclinical models. Male mice treated with the non-selective ROCK inhibitor fasudil, had increased CBF values at baseline compared to control mice, however there were no differences in regional CBF between groups following transient MCAo (Rikitake, et al., 2005). Alternatively, non-selective ROCK inhibition did attenuate the CBF deficit in mice subjected to distal MCAo, but not in mice lacking the gene for eNOS (*eNOS*^*-/-*^), suggesting an endothelial-dependent mechanism of neuroprotection (Shin, et al., 2007). On the other hand, selective inhibition of the ROCK2 isoform with KD025 did not improve absolute CBF values of male mice following distal MCAo, but did reduce the overall area of tissue with a severe perfusion deficit (Lee, et al., 2014).

Finally, heterozygous knockout of ROCK2, but not ROCK1, in male mice improved absolute CBF following 1-hour of transient MCAo as measured by an indicator fractionation technique (Hiroi, et al., 2018). While there is some evidence that ROCK inhibition and deletion improves CBF outcomes following stroke, this was not observed in the current study. This may be due to the nature of the stroke model used, wherein PT stroke does not provide reperfusion injury and is less sensitive to measurements of neuroprotection. A failure to detect CBF improvement could also be due to the technique used to measure CBF. In studies that do show some improvement in CBF following stroke, the techniques used offer a larger picture of total CBF deficit in the entire brain, whereas LDF provides high temporal resolution of only a very small area surrounding the infarct core. It is possible that there may have been CBF improvements in *ROCK2*^*+/-*^ mice in the current study as has been previously described in other studies (Hiroi, et al., 2018; Lee, et al., 2014), but simply could not be detected with the LDF method used.

Haploinsufficiency for the ROCK2 gene in intact female mice demonstrated profound differences in CBF values following PT stroke. Compared to Intact WT females, Intact *ROCK2*^*+/-*^ females had significantly higher CBF values in the immediate 30-minute time period following stroke. Additionally, Intact *ROCK2*^*+/-*^ females showed the same delayed drop in CBF values that was observed in both groups of intact males, as well as in Ovx WT females, wherein the maximal drop in CBF was observed at 48 hours following stroke. This sex-specific role of ROCK2 in CBF outcomes may be due to differences in RhoA/ROCK signaling in platelets, which has been shown to be upregulated in platelets from females, corresponding to increased levels of phosphorylated MLC and platelet hyperreactivity in females (Schubert, et al., 2016). It is possible that increased RhoA/ROCK signaling in platelets makes them more reactive to thrombus formation during PT stroke induction, therefore reducing the amount of ROCK2 available in hyperreactive female platelets could increase the necessary time for thrombosis to occur, resulting in the phenotypic male-like CBF response observed in this particular model. Indeed, platelets from male mice that harbor a ROCK2 deletion selectively in platelets were shown to be less prone to aggregation and thrombosis, and also had improved CBF following a thromboembolic model of MCAo compared to WT mice (Sladojevic, et al., 2017), further supporting this hypothesis.

### Interactions Between Sex Hormones and ROCK2 in CBF Outcomes Following PT Stroke

Contrary to comparisons made between males and females in intact WT mice, there were no differences in the behaviour of CBF at any timepoint post stroke between sexes in intact *ROCK2*^*+/-*^ mice. Removal of endogenous sex hormones from *ROCK2*^*+/-*^ males and females also did not alter the CBF response at any timepoint post stroke. Overall, regardless of genotype or gonadectomy status, male mice showed a phenotypic response in CBF following stroke where they do not reach a maximal drop in CBF that is expected with stroke until 48 hours following stroke induction. Although not statistically significant from the measurements taken immediately post stroke, Gdx *ROCK2*^*+/-*^ males still displayed peak drop in CBF values when measured at 48 hours following stroke induction. This may be due in part to the removal of endogenous testosterone. Low testosterone observed with aging and vascular disease in males is associated with the upregulation of ROCK (Sopko, et al., 2014). Consistent with the current hypothesis, if upregulation of ROCK in platelets promotes quicker aggregation of platelets, this may account for slightly lower CBF values immediately following stroke in both WT and ROCK2+/- males, however the upregulation of ROCK may not be large enough to produce the same response as seen in females. Meanwhile, removal of endogenous sex hormones from WT females produced the same phenotype as observed in males. Lastly, *ROCK2*^*+/-*^ females also showed the phenotypic response of delayed CBF drop, which was unaffected by removing endogenous sex hormones.

Interestingly, there is some evidence that estrogen may increase ROCK activity in females. Superior mesenteric arteries isolated from rats showed greater constriction in response to norepinephrine in young adult female rats aged 8-24 weeks, but not in adolescent (4 week) or aged (1 and 1.5 years) female rats. This increased vascular reactivity correlated with increased estrogen levels in young adult females (Li, et al., 2014). Furthermore, protein levels of ROCK were significantly greater in mesenteric arteries from both young adult and aged female rats compared to age-matched males. Short-term incubation with estrogen increased ROCK activity in mesenteric arteries, and long-term incubation increased protein levels of ROCK, suggesting both genomic and non-genomic regulation of ROCK by estrogen (Li, et al., 2014). Given the potential for estrogen to upregulate RhoA/ROCK signaling, it is possible that removing endogenous estrogens downregulates ROCK and therefore reduces its activity in platelets, reducing thrombus formation in this particular model of ischemic stroke. Indeed, as described earlier, estrogen treatment may also contribute to increased reactivity and aggregation of platelets. Therefore, decreased estrogen may alter RhoA/ROCK signaling in platelets, contributing to a decrease in their reactivity and potentially delaying aggregation and thrombus formation during PT stroke induction.

### Infarct Volumes and Nitric Oxide Staining

There were no differences in stroke lesion volumes as determined by MRI measured 48 hours following stroke induction. Although previous studies have reported a neuroprotective role of ROCK inhibition or deletion in ischemic stroke and infarct volume reduction, these were all reported using variations of the MCAo model. While MCAo is useful for characterizing neuroprotection, the model produces quite large strokes in which up to half of the affected hemisphere is lesioned, which has been argued to not be representative of clinical strokes seen in humans (Carmichael, 2005). This study is the first to characterize ROCK2 deletion in a small, focal ischemic stroke, that may better mimic clinical conditions, and which may be less sensitive to reductions in infarct size when the infarct size is not as large to begin with. Importantly, because there were no differences in infarct volume between groups, it confirms that differences observed in CBF measurements are not due to a larger area of injury which correlates with a more severe CBF deficit.

ROCK negatively regulates eNOS function, providing speculation that the neuroprotective role of ROCK inhibition and deletion is mediated through upregulation of eNOS. Given the evidence that *ROCK2*^*+/-*^ mice have increased eNOS levels in brain ECs (Hiroi, et al., 2018) we assessed levels of NO in the infarct 24 hours following PT stroke induction using DAF-FM diacetate staining. There were no differences in NO levels in any group following PT stroke, despite a previous study using the same stain showing increased NO levels at the same timepoint following transient MCAo in *ROCK2*^*+/-*^ male mice compared to both *ROCK1*^*+/-*^ and WT mice (Hiroi, et al., 2018). Again, this may be due to differences in mechanisms of ischemic stroke models. Unlike MCAo, the PT stroke model does not have any reperfusion injury, so eNOS upregulation may be more prevalent in stroke models with reperfusion. If this were true, haploinsufficiency of ROCK2 would reduce eNOS inhibition resulting in increased NO bioavailability following MCAo. Additionally, NO has an extremely short half life and staining may not have been done quickly enough to detect any differences. Further experiments such as eNOS and ROCK activity assays would be required to solidify the absence of eNOS upregulation in *ROCK2*^*+/-*^ mice following PT stroke in the somatosensory cortex.

### Limitations and Future Directions

To support the hypothesis that Intact WT females have a more severe reduction in CBF following PT stroke due to accelerated thrombosis, further experimentation would be required to support this. Performing a coagulation assay would first be necessary to assess if Intact WT females are more prone to blood coagulation than other groups. From there, assessing platelet reactivity and also determining ROCK activity in platelets in these groups would further elucidate the precise mechanisms involved. It would also be interesting to assess eNOS activity and ROCK activity in brain ECs to determine if these are upregulated following PT stroke, as most studies characterizing these have been performed using MCAo.

Additionally, all CBF measurements were performed under K/X anesthesia. This anesthesia was chosen because it has lesser effects on vasodilation than other vaporized anesthetics such as isoflurane (Rakymzhan, et al., 2021), which also has neuroprotective properties against ischemic stroke via modulation of eNOS (Lu, et al., 2017; Krolikowski, et al., 2006; Kehl, et al., 2002). However, there is still evidence that ketamine can cause vasodilation and alter CBF (Rakymzhan, et al., 2021). Ideally, these experiments would be repeated in awake and unanesthetized mice, which has been successfully performed with PT stroke using cranial window preparations in rats (Yang, et al., 2019; Lu, et al., 2014) and mice (He, et al., 2020). This would better mimic clinical stroke conditions and eliminate any confounding effects of anesthetics.

## Conclusion

Overall, there is a clear sex difference in CBF outcomes following stroke in this model of PT stroke in the somatosensory cortex, in which ROCK2 appears to be implicated. All groups except for Intact WT females show a delayed drop in CBF values, reaching a maximal drop in CBF at 48 hours following stroke induction. This is likely due to mechanistic differences in stroke induction, where both ROCK2 and endogenous female sex hormones, most likely estrogen, mediate platelet aggregation and thrombus formation resulting in delayed obstruction of the vessels in the infarct. Sex differences in response to different stroke models reiterates the importance of studying both sexes in the pathogenesis of disease. Characterization of stroke models in both males and females is of the utmost importance in preclinical research to prevent translational failure. This research helps clarify the role of ROCK2 in CBF outcomes following a PT stroke, and brings to light new questions about interactions between rho-kinase and sex hormones in the cerebrovasculature.

## Acknowledgements

We would like to thank the University of Ottawa’s animal care and surgical facilities for providing the space and expertise required to successfully perform in vivo experiments. *ROCK2*^*+/-*^ mice were a generous gift from Dr. Zheng Ping Jia (University of Toronto, Canada).

